# Fingertip real contact area scales quadratically with input voltage in electrostatic actuation

**DOI:** 10.64898/2026.05.31.729125

**Authors:** Celal Umut Kenanoglu, Yasemin Vardar

**Affiliations:** Department of Cognitive Robotics, Delft University of Technology, Delft, 2628 CD, The Netherlands

**Keywords:** Electrovibration, Electroadhesion, Contact mechanics

## Abstract

Touchscreens have become the dominant interface in consumer electronics, yet interactions with them remain primarily visual. Incorporating haptic feedback that simulates touch sensations could make these interactions more natural and intuitive. Electrostatic actuation, which modulates friction by attracting the finger toward a capacitive surface using an alternating voltage, offers a promising approach. The resulting increase in friction is often attributed to the rise in real contact area; however, direct experimental evidence linking voltage input parameters to real contact area and contact forces remains limited. Here, we use frustrated total internal reflection to directly image the real contact area while simultaneously measuring contact forces during controlled finger sliding under electrostatic actuation. We systematically vary voltage amplitude (75–150 V) and excitation frequency (30–230 Hz) and quantify the changes in contact area and forces as functions of these parameters. Our results reveal a quadratic dependence of real contact area, electrostatic attraction, and tangential force on voltage amplitude, with comparatively small effects of excitation frequency. These findings clarify the respective roles of voltage amplitude and frequency in the electrostatic modulation of finger contact mechanics, providing design guidelines for haptic display design.

## 1 Introduction

Touchscreens are now predominant interface for personal electronic devices and public terminals; however, interaction with these systems remains largely dominated by visual and auditory modalities. The absence or ineffectiveness of tactile feedback makes interaction less natural and, consequently, less intuitive. Surface haptics aims to overcome this limitation by enabling programmable tactile cues, such as friction and texture, on flat surfaces. Among surface haptics technologies, electrostatic actuation has been increasingly adopted in research labs due to its relative simplicity and ease of integration. In this approach, an alternating voltage is applied to the touchscreen’s conductive layer, generating a periodic attractive force between the surface and the fingertip [5]. Temporal modulation of this voltage allows control of interfacial friction, producing tactile sensations [5, 7, 13, 22]. Despite the maturity of this technology, its broader adoption remains limited, partly due to an incomplete understanding of the coupling between actuation and fingertip mechanics, which hinders precise and predictable control of the resulting tactile effects [17].

During electrostatic actuation, the fingertip skin is alternately attracted toward and released from the surface. Numerous studies have reported an increase in tangential force during this process [6,14,18,19,23]. This increase is commonly attributed to the additional electrostatic attraction at the interface, whose response has been shown to depend on excitation frequency, the applied voltage, and the coupled electrical, mechanical, and geometrical properties of the finger– touchscreen interface [1, 16, 18, 19]. This interaction is typically modeled using a parallel-plate capacitance model [25]. Under this model, the electrostatic force scales quadratically with the applied voltage, a relationship that has been experimentally confirmed through measurements of contact forces between a sliding finger and an electrostatic display [6, 14, 24]. However, force measurements alone do not directly reveal how electrostatic actuation alters the underlying contact mechanics during sliding, such as changes in the real contact area. According to classical friction theory, rise in tangential force is often associated with increase in real contact area [9]. Despite this observation, direct experimental measurements of how real contact area varies jointly with voltage amplitude and excitation frequency during electrostatic actuation remain limited.

Although limited, a small number of studies have examined the changes in finger contact area under electrostatic actuation using optical measurements or computational approaches. Sirin *et al*. experimentally investigated fingerpad contact under electrostatic actuation and surprisingly reported a decrease in apparent contact area (i.e., fingerpad boundary) when actuation was applied [21]. They attributed the observed increase in tangential force to an increase in real contact area; however, the real contact area was not directly measured. In parallel, several studies have interpreted friction increases under electrostatic actuation using contact-mechanics models, estimating actuation-induced increases in real contact area through mean-field, multiscale approaches [2, 3, 15, 20]. These studies predicted a rise in real contact area as a function of voltage and excitation frequency, and validated the models indirectly using tangential-force measurements. To the authors’ knowledge, [12] is the only study that experimentally demonstrated an increase in real contact area under electrostatic actuation; however, the experiments were conducted at a fixed voltage. Consequently, direct experimental evidence that simultaneously quantifies real contact area, electrostatic attraction, and tangential force while jointly varying voltage amplitude and excitation frequency during sliding is still lacking.

Here, we address this gap by simultaneously measuring real contact area and contact forces during controlled finger sliding while systematically varying voltage amplitude (75–150 V) and excitation frequency (30–230 Hz). We demonstrate that voltage is the dominant driver of changes in real contact area, electrostatic attraction, and tangential force. Finally, we provide compact quadratic fits for all three measures, consistent with the voltage-squared dependence predicted by parallel-plate electrostatic models, providing a low-parameter basis for modeling electrostatic surface haptics.

## 2 Methods

### 2.1 Electrostatic Actuation

Electrostatic actuation modulates friction by generating an oscillating electric field that attracts the skin toward the surface. The resulting electrostatic force *F*_*e*_ depends on the input voltage, the effective gap area, the gap thickness between the dielectric layers, and the dielectric properties of the layers separating the conductive bodies, and can be calculated from parallel-plate capacitance theory, as follows [18, 19, 25]:

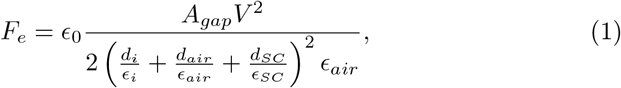

where *A*_*gap*_ is the effective gap area at the fingertip–surface interface, *V* is the input voltage. *d*_*i*_, *d*_*SC*_, and *d*_*air*_ represent the thicknesses of the insulating layer of the touchscreen, stratum corneum, and the air gap between the touchscreen and finger, respectively. These layers act as dielectric and can be treated as a series stack that sets an effective electrical separation between the two conductive layers. Their relative permittivity are denoted as *ϵ*_*i*_, *ϵ*_*SC*_, and *ϵ*_*air*_, while *ϵ*_0_ is the electric permittivity of the vacuum; see Fig. 1a for a schematic of the finger-surface interface.

**Fig. 1:**
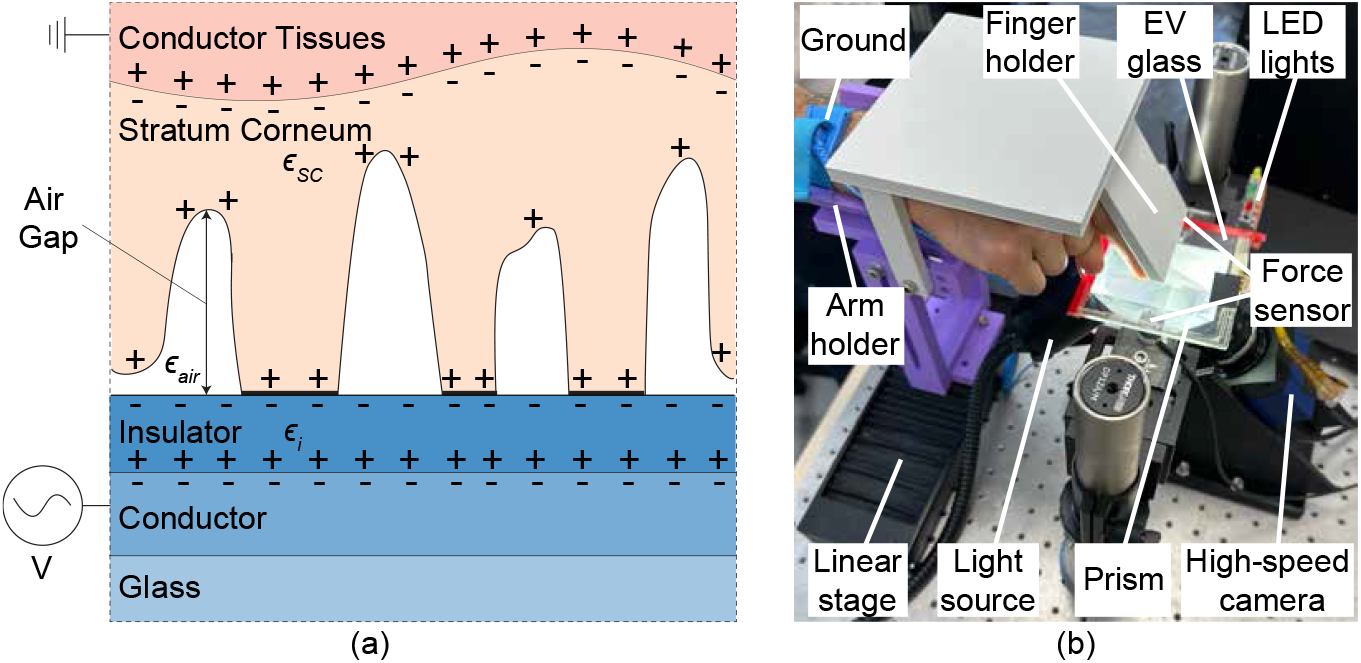
(a) Schematic of the multilayer fingertip–surface interface under an applied AC voltage. Here, *ϵ*_*i*_, *ϵ*_*SC*_, and *ϵ*_*air*_ denote the relative permittivity of the touchscreen insulating layer, the stratum corneum, and the air gap between the touchscreen and the fingertip, respectively. Opposite charges form at the interface and alternate with the voltage polarity. (b) Experimental setup combining six-axis force sensing and FTIR-based imaging to measure contact forces and real contact area during sliding.

### 2.2 Experimental Procedure

During experiments, we visualized changes in finger contact area using frustrated total internal reflection (FTIR) imaging [8] while simultaneously measuring contact forces as one participant slid their finger on a capacitive touchscreen; see Fig. 1b for the experimental setup. The touchscreen (SCT3250, 3M Inc.) was mounted on two six-axis force sensors (Nano17 Titanium, ATI Inc.). It was actuated by a data acquisition card (SCB-68A, NI Inc.) with a sampling rate of 10 kHz and amplified with a high-voltage amplifier (9200A, Tabor Electronics). Another data acquisition card (PCIe 6321, NI Inc.) was used to record contact forces at the same sampling rate. Fingertip contact area was imaged from below the glass using a high-speed camera (MotionBLITZ EoSens mini2, Mikrotron) with a lens (LM16HC, Kowa) at 1000 frame per second (fps). Finger motion was imposed by a motorized linear stage (NRT150/M, Thorlabs Inc.), with the finger contacting the surface at a fixed angle of 60°. Participant regulated the normal force using real-time LED feedback to maintain the force within *±*10% of the target. The glass was illuminated by a light source (KL 2500 LED, Schott) equipped with a custom collimation package and diffuser. A fan was used to maintain consistent skin dryness during sliding.

The study adhered to the Declaration of Helsinki and was approved by the Ethics Council of TU Delft (application no. 5108). The participant provided informed consent. Before the experiment, the participant washed their hands and dried them with a microfiber cloth. The experiment consisted of five sessions, each conducted on a separate occasion. In each session, all combinations of four input voltage amplitudes (75 V, 100 V, 125 V, and 150 V) and five sine-wave frequencies (30 Hz, 80 Hz, 130 Hz, 180 Hz, and 230 Hz) were tested, with three repetitions per frequency–amplitude combination (4 amplitudes × 5 frequencies × 5 sessions × 3 repetitions = 300 trials). Each trial consisted of a voltage-off phase followed by a voltage-on phase within the same sliding stroke. In each trial, the fingertip was moved laterally at a constant speed of 20 mm/s using the motorized stage, while the participant regulated their normal force to 1 N using real-time visual feedback. Data were retained only when the measured normal force remained within *±*10% of the target and the fingerprint image was visible. For each frequency–voltage combination, results were computed by averaging across the three repetitions within that session. Each session lasted approximately 30 minutes.

Data acquisition from the force sensors, linear stage control, and camera triggering was synchronized in MATLAB/Simulink. All signals other than contact images were generated and recorded in Simulink, whereas images were acquired using MotionBLITZ software. A trigger was issued from Simulink when the finger entered the camera field of view; only the normal force (*F*_*n*_) and tangential force (*F*_*t*_) data within this window were used for analysis. Before area extrac-tion, raw FTIR images were geometrically corrected. Radial lens distortion was removed using intrinsic parameters from checkerboard calibration, and a projective transformation was applied to obtain an equivalent top-down view [10]. The homography was calibrated by mapping the imaged ellipse of a circular rubber target placed on the glass to a true circle. We used FTIR imaging to estimate the real contact area, defined here as the optically resolved portion of the interface in intimate contact (i.e., the sum of micro-contact regions). This estimation was computed from the corrected images following [11]. However, as the real contact area is inherently scale-dependent [15, 16], the FTIR-based estimate may differ from the true contact area. For each trial, we computed the real contact area ratio *A*^on^*/A*^off^, where *A*^on^ and *A*^off^ denote the mean real contact area during the voltage-on and voltage-off phases within the same sliding stroke. Using the same within-stroke segmentation, we also computed the tangential-force ratio 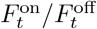 and estimated the electrostatic attraction force as *F*_*e*_ = (1 − *µ*_off_ */µ*_on_)*F*_*n*_, where *µ* = *F*_*t*_*/F*_*n*_ is the friction coefficient [2, 6]. For statistical analysis, session-level means were used in the two-way ANOVA. Across all trials, the mean normal force remained similar between the voltage-OFF and voltage-ON conditions (1.006 *±* 0.018 N vs 1.02 *±* 0.019 N).

## 3 Results and Discussion

Our results show that electrostatic actuation increases real contact area primarily with voltage amplitude, with comparatively smaller frequency effects (Figs. 2a and 2b). As voltage increases, the dark regions in the FTIR images become more pronounced, indicating an increase in real contact area (Fig. 2a and Supplementary Movies). Consistently, the normalized area ratio *A*^on^*/A*^off^ increases monotonically with voltage across all tested frequencies (Fig. 2b). This voltage-driven increase in real contact area can be attributed to the additional electrostatic attraction in the normal direction acting at the interface, which increases the effective normal loading and, in turn, the tangential force [6, 19, 21, 23]. Consistent with Eq. 1, a higher voltage produces a larger attractive force, which can promote additional micro-junction formation and increase the real contact area. Across frequencies, the curves show only modest shifts, with the largest modulation at intermediate frequencies. The reduced modulation at 30 Hz may reflect the frequency-dependent electrical response of the fingertip–surface interface, characterized by increased charge leakage at low frequencies [2, 20]. The reduced modulation at 230 Hz is consistent with the viscoelastic behavior of the fingertip, which may limit its ability to follow rapid actuation at higher frequencies [4, 12]. Similar U-shaped behavior has also been reported in previous studies [1, 2, 12].

**Fig. 2:**
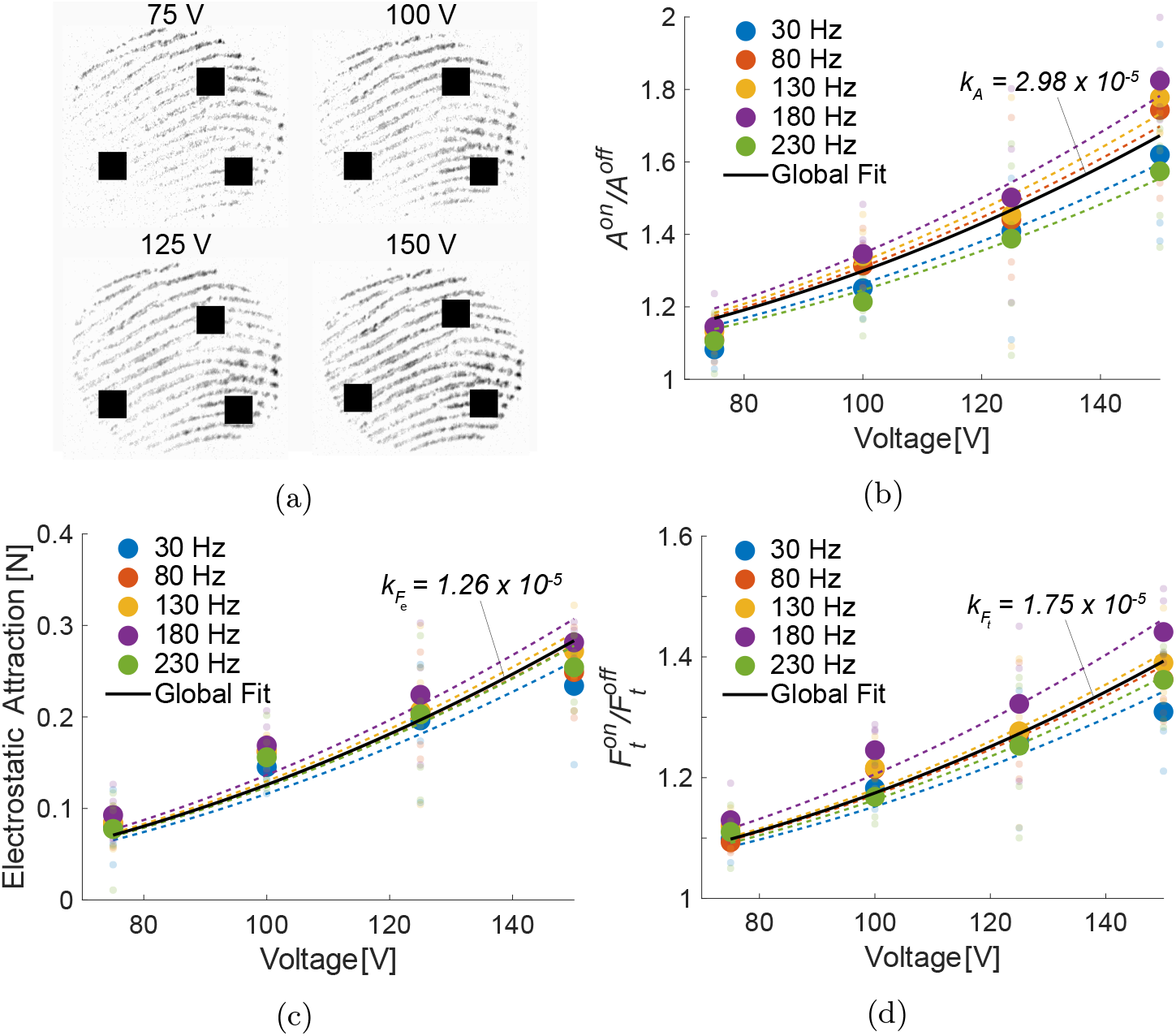
Experimental results. (a) Representative fingertip contact images at different input voltage amplitudes. (b) Real contact area ratio *A*^on^*/A*^off^, (c) estimated electrostatic attraction force *F*_*e*_, and (d) tangential force ratio 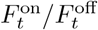 plotted versus voltage amplitude for each excitation frequency. Solid black lines show frequency-independent global fits 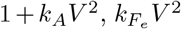, and 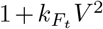 for each metric, respectively; dashed lines indicate frequency-specific fits. Large circles denote session means; smaller shaded circles represent repetition means within each session. Across the five sessions, the median 95% confidence-interval half-width was 0.133 for *A*^*on*^*/A*^*on*^, 0.040 N for *F*_*e*_, and 0.069 for 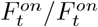.

Finally, we observed that the real contact area of the fingertip and tangential force oscillates at twice the frequency of the input voltage (Fig. 3 and Supplementary Movies), consistent with electrostatic force behavior and in agreement with prior reports [12, 23, 24]. Representative traces show that modulation of tangential force and real contact area primarily occurs during the voltage-ON phase, while the normal force remains within the *±*10% band around 1 N.

**Fig. 3:**
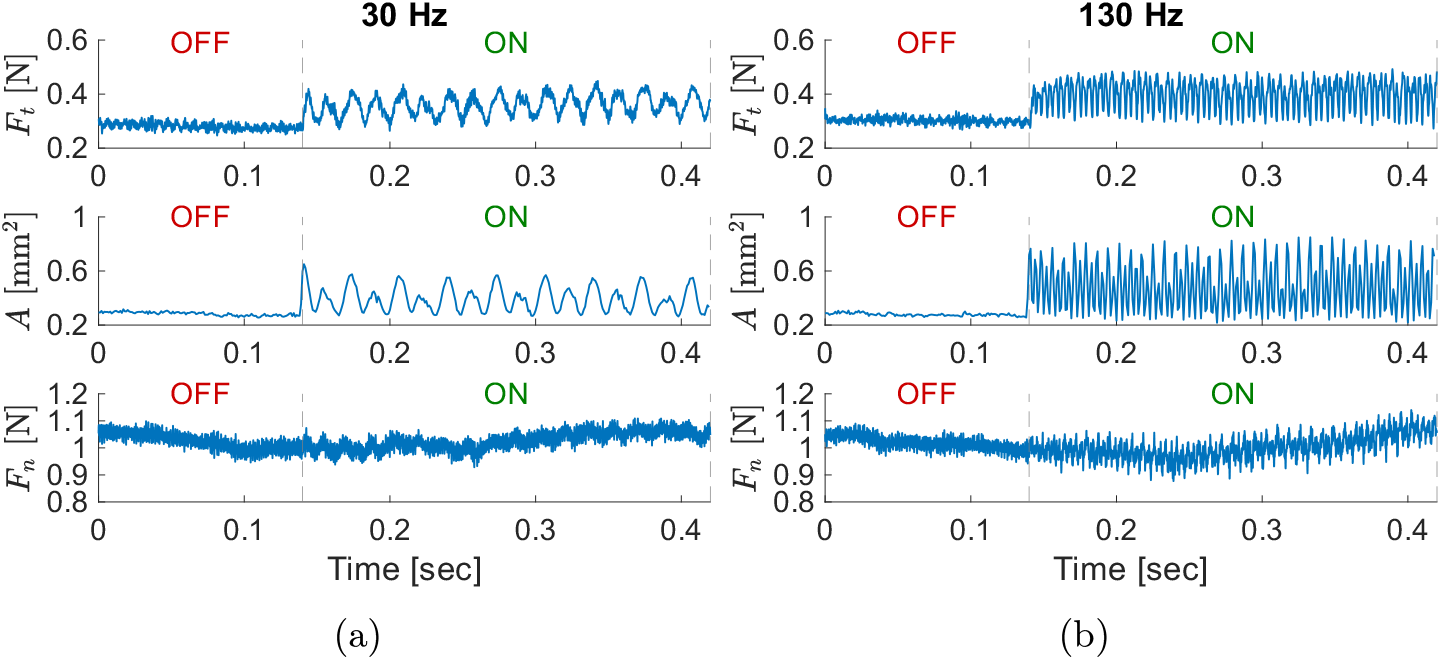
Time-domain response under electrostatic actuation. Representative traces of tangential force *F*_*t*_, real contact area *A*, and normal force *F*_*n*_ during sliding under electrostatic actuation at (a) 30 Hz and (b) 130 Hz. The dashed line indicates the transition from the voltage-OFF to the voltage-ON condition.

To quantify the dependence of real contact area to voltage amplitude and excitation frequency, we analyzed *A*^on^*/A*^off^ using a two-way ANOVA with main factors of amplitude (75–150 V) and frequency (30–230 Hz). Before hypothesis testing, we assessed residual normality using the Lilliefors test, which did not reject the null hypothesis of normality. The analysis showed a strong main effect of amplitude (*F* (3, 80) = 65.54, *p* = 1.71 × 10^*−*21^) and a smaller main effect of frequency (*F* (4, 80) = 2.67, *p* = 0.0381), with no amplitude × frequency interaction (*F* (12, 80) = 0.304, *p* = 0.987). Bonferroni-corrected post-hoc com-parisons confirmed that the results across all voltage amplitude levels differed significantly from one another (*p <* 0.001), whereas pairwise comparisons across frequencies were not significant. Together, these results indicate that voltage amplitude is the dominant driver of contact-area modulation in our experiments, while frequency effects are comparatively small for the tested ranges.

Given the absence of a statistically significant interaction effect, the comparatively small main effect of frequency, and the lack of significant pairwise differences across frequencies, we used a frequency-independent global fit to capture the dominant voltage dependence (Fig. 2b; frequency-specific fits are shown as dashed lines). Motivated by the quadratic scaling of electrostatic attraction in Eq. 1, we modeled the real area ratio as *A*^*on*^*/A*^*off*^ = 1 + *k*_*A*_*V* ^2^. Fitting the pooled data to this model yielded *k*_*A*_ = 2.98 × 10^*−*5^ V^-2^, with a root mean square error (RMSE) of = 0.154 and a mean absolute error (MAE) of = 0.113.

Our findings demonstrate that the estimated electrostatic attraction force also increases monotonically with voltage amplitude (Fig. 2c), in agreement with prior work [6, 14, 24]. To further quantify its dependency on voltage amplitude and frequency with a two-way ANOVA after verifying that the normality of residuals was not rejected by the Lilliefors test. The analysis revealed a strong effect of amplitude (*F* (3, 80) = 79.62, *p* = 5.97 × 10^*−*24^), but no effect of frequency (*F* (4, 80) = 1.25, *p* = 0.2968) and no significant interaction effect (*F* (12, 80) = 0.131, *p* = 0.9998). Bonferroni-corrected post-hoc comparisons confirmed that the results across all amplitude levels differed significantly (*p <* 0.001).

To capture the dependence of the electrostatic attraction force on voltage amplitude, we fitted the model 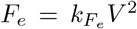 to the pooled data, yielding 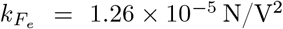 with RMSE of = 0.0446 N and MAE of = 0.0352 N.

This quadratic scaling is consistent with prior findings reporting a similar de-pendence of electrostatic force to voltage amplitude [24], based on experiments with 15 participants (coefficient 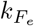 was found as 1.34 × 10^*−*5^ N/V^2^).

We further compared the experimentally obtained coefficient 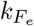 with the theoretically calculated value from Eq. 1 [6, 18]. For this calculation, we adopted representative literature values for the layer thicknesses *d*_i_, *d*_SC_, as well as the permittivity values *ϵ*_i_, *ϵ*_SC_, and *ϵ*_air_, namely 1 *µ*m, 200 *µ*m, 3.9, 3000, and 1, respectively [2, 6]. The permittivity of the interfacial air-gap layer was approximated as that of air as a simplifying assumption, although the actual interfacial medium may deviate from pure air due to moisture and other interface-specific effects. The interfacial air-gap thickness *d*_air_ was treated as an effective fitted parameter (*d*_air_ = 8.47 *µ*m), since it is expected to depend strongly on the contact condition, fingertip roughness, and related interface-specific factors. This fitted value is of the same order of magnitude as the average interfacial separation reported in [15]. For the area term in Eq. 1, we used the apparent contact area as an approximation of the effective gap area, following prior work [6, 18]. This approximation is reasonable because, in rough contacts, the real contact area is generally only a small fraction of the apparent area [6, 16]. Consistently, in our measurements, the FTIR-estimated real contact area was only about 1% of the apparent fingertip area, such that the gap area was dominated by the non-contact portion of the interface. From the FTIR images, we estimated an apparent contact area (i.e., the fingertip boundary area) of 55 mm^2^. Using these values, the predicted coefficient is 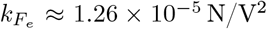, in close agreement with the fitted value.

Moreover, tangential force increased with voltage across all tested frequencies (Fig. 2d), consistent with the trends in real area and electrostatic attraction. To assess the effects of voltage amplitude and frequency, we first verified that residual normality was not rejected by the Lilliefors test. We then conducted a two-way ANOVA with factors amplitude and frequency. The analysis revealed a strong effect of amplitude (*F* (3, 80) = 75.46, *p* = 2.93 × 10^*−*23^) and a smaller effect of frequency (*F* (4, 80) = 3.52, *p* = 0.0106), with no significant amplitude×frequency interaction (*F* (12, 80) = 0.453, *p* = 0.9355). Bonferroni-corrected post-hoc comparisons confirmed that the results across all amplitude levels differed significantly from each other (*p <* 0.001). For the results across frequencies, significant differences were observed only between 30 Hz and 180 Hz (*p* = 0.009) and 180 Hz and 230 Hz (*p* = 0.043). Given the absence of an interaction and the comparatively small main effect of frequency, we fitted a frequency-independent quadratic model, 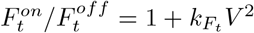, yielding 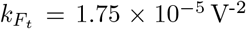 with RMSE of = 0.0673 and MAE of = 0.0511. Overall, the tangential force increase is consistent with increases in real area and electrostatic attraction reported above.

## 4 Conclusion

This study experimentally quantified how electrostatic actuation modulates key aspects of fingertip contact mechanics during sliding—real contact area, tangen-tial force, and estimated electrostatic attraction—as a function of voltage amplitude and excitation frequency. Across all tested excitation frequencies (30–230 Hz), increasing voltage amplitude consistently increased the real contact area, the estimated electrostatic attraction, and the tangential force. For all three measures, voltage amplitude was the dominant factor, with comparatively weak effects of frequency and no observable interaction between the two. These results support a concise description of electrostatic modulation in which changes in real contact area and tangential force primarily scale with the square of the voltage, consistent with the expected voltage dependence of electrostatic attraction.

The present results are based on a single participant, a single sliding condition, and a limited voltage—frequency range. Accordingly, these findings should be interpreted within this context. As fingertip contact is known to vary with skin hydration, individual properties, and operating conditions [12], future work will extend the study to multiple participants and a broader range of forces and speeds, and will investigate how inter-individual variability and environmental factors affect the actuation, enabling broader generalization.

## Acknowledgments

This work has been partially supported by the Dutch Research Council, NWO, with the project number 20624.

## Disclosure of Interests

The authors declare no conflict of interest.

## Author contributions

CUK: Conceptualization, Methodology, Investigation, Software, Hardware, Formal Analysis, Data Curation, Visualization, Writing - original draft. YV: Conceptualization, Methodology, Formal Analysis, Visualization, Writing - original draft, Writing - review & editing, Supervision, Resources, Funding Acquisition.

## Data availability

The dataset used in this study is publicly available via 4TU.ResearchData under the following DOI: 10.4121/08aa3d2d-1761-4e67-a993-0eff1c870c37.

## Notes

### Competing Interest Statement

The authors have declared no competing interest.

## References

1. AliAbbasi, E., Martinsen, Ø.G., Pettersen, F.J., Colgate, J.E., Basdogan, C.: Experimental estimation of gap thickness and electrostatic forces between contacting surfaces under electroadhesion. Advanced Intelligent Systems 6(4), 2300618 (2024)

2. AliAbbasi, E., Sormoli, M.A., Basdogan, C.: Frequency-dependent behavior of electrostatic forces between human finger and touch screen under electroadhesion. IEEE Transactions on Haptics 15(2), 416–428 (2022)

3. Ayyildiz, M., Scaraggi, M., Sirin, O., Basdogan, C., Persson, B.N.: Contact mechanics between the human finger and a touchscreen under electroadhesion. Proceedings of the National Academy of Sciences 115(50), 12668–12673 (2018)

4. Balasubramanian, J.K., Pool, D.M., Vardar, Y.: Sliding speed influences electrovibration-induced finger friction dynamics on touchscreens. Tribology International 213(11105), 4 (2026)

5. Basdogan, C., Giraud, F., Levesque, V., Choi, S.: A review of surface haptics: Enabling tactile effects on touch surfaces. IEEE Transactions on Haptics 13(3), 450–470 (2020)

6. Basdogan, C., Sormoli, M.R.A., Sirin, O.: Modeling sliding friction between human finger and touchscreen under electroadhesion. IEEE Transactions on Haptics 13(3), 511–521 (2020)

7. Bau, O., Poupyrev, I., Israr, A., Harrison, C.: Teslatouch: electrovibration for touch surfaces. In: Proceedings of the 23nd Annual ACM Symposium on User Interface Software and Technology. pp. 283–292 (2010)

8. Bochereau, S., Dzidek, B., Adams, M., Hayward, V.: Characterizing and imaging gross and real finger contacts under dynamic loading. IEEE Transactions on Haptics 10(4), 456–465 (2017)

9. Bowden, F.P., Tabor, D.: The friction and lubrication of solids. Oxford University Press (2001)

10. Goshtasby, A.: Piecewise linear mapping functions for image registration. Pattern Recognition 19(6), 459–466 (1986)

11. Huloux, N., Willemet, L., Wiertlewski, M.: How to measure the area of real contact of skin on glass. IEEE Transactions on Haptics 14(2), 235–241 (2021)

12. Kenanoglu, C.U., Wiertlewski, M., Vardar, Y.: Electrostatic actuation induces competing adhesion and vibration regimes at fingertip contact. Advanced Intelligent Systems, e70433 (2026)

13. Mallinckrodt, E., Hughes, A., Sleator Jr, W.: Perception by the skin of electrically induced vibrations. Science 118(3062), 277–278 (1953)

14. Meyer, D.J., Peshkin, M.A., Colgate, J.E.: Fingertip friction modulation due to electrostatic attraction. In: 2013 World Haptics Conference (WHC). pp. 43–48. IEEE (2013)

15. Persson, B.: The dependency of adhesion and friction on electrostatic attraction. The Journal of chemical physics 148(14) (2018)

16. Persson, B.N.: General theory of electroadhesion. Journal of Physics: Condensed Matter 33(43), 435001 (2021)

17. Pool, D.M., Vardar, Y.: Fits: Ensuring safe and effective touchscreen use in moving vehicles. IFAC-PapersOnLine 58(30), 248–253 (2024)

18. Shultz, C., Peshkin, M., Colgate, J.E.: The application of tactile, audible, and ultrasonic forces to human fingertips using broadband electroadhesion. IEEE Transactions on Haptics 11(2), 279–290 (2018)

19. Shultz, C.D., Peshkin, M.A., Colgate, J.E.: Surface haptics via electroadhesion: Expanding electrovibration with johnsen and rahbek. In: 2015 World Haptics Conference (WHC). pp. 57–62. IEEE (2015)

20. Sirin, O., Ayyildiz, M., Persson, B.N., Basdogan, C.: Electroadhesion with application to touchscreens. Soft matter 15(8), 1758–1775 (2019)

21. Sirin, O., Barrea, A., Lefèvre, P., Thonnard, J.L., Basdogan, C.: Fingerpad contact evolution under electrovibration. Journal of the Royal Society Interface 16(156), 20190166 (2019)

22. Vardar, Y.: Tactile perception by electrovibration. Springer (2020)

23. Vardar, Y., Güçlü, B., Basdogan, C.: Effect of waveform on tactile perception by electrovibration displayed on touch screens. IEEE Transactions on Haptics 10(4), 488–499 (2017)

24. Vardar, Y., Kuchenbecker, K.J.: Finger motion and contact by a second finger influence the tactile perception of electrovibration. Journal of the Royal Society Interface 18(176), 20200783 (2021)

25. Vodlak, T., Vidrih, Z., Vezzoli, E., Lemaire-Semail, B., Peric, D.: Multi-physics modelling and experimental validation of electrovibration based haptic devices. Biotribology 8, 12–25 (2016)

